# *Staphylococcus epidermidis* metabolic adaptation and biofilm formation in response to varying oxygen

**DOI:** 10.1101/665356

**Authors:** Ulrik H. Pedroza-Dávila, Cristina Uribe-Alvarez, Lilia Morales-García, Emilio Espinoza-Simón, Adriana Muhlia-Almazán, Natalia Chiquete-Félix, Salvador Uribe-Carvajal

## Abstract

*Staphylococcus epidermidis* is a Gram-positive saprophytic bacterium found in the microaerobic/anaerobic layers of the skin. It becomes a health hazard when introduced across the skin by punctures or wounds. *S. epidermidis* forms biofilms in low O_2_ environments. As oxygen concentrations ([O_2_]) decreased, the metabolism of *S. epidermidis* was modified ranging from fully aerobic to anaerobic. Respiratory activity increased at high [O_2_], while anaerobically grown cells exhibited the highest rate of fermentation. High aerobic metabolism coincided with high hydrogen peroxide-mediated damage. Remarkably, the rate of growth decreased at low [O_2_] even though the concentration of ATP was high. Under these conditions bacteria associated into biofilms. Then, in the presence of metabolic inhibitors, biofilm formation decreased. It is suggested that when [O_2_] is low *S. epidermidis* accumulates ATP in order to synthesize the proteins and polysaccharides needed to attach to surfaces and form biofilms.

**Importance:** Bacteria and humans coexist, establishing all kinds of relationships that may change from saprophytic to infectious as environmental conditions vary. S. epidermidis is saprophytic when living in the skin. Inside the organism it evokes a pathologic reaction and is thus rejected by the organism. Additionally it is forced to adapt to high oxygen concentrations, becoming vulnerable to reactive oxygen species, which may come from leukocyte attack. Avoiding both, high oxygen and leukocytes is a must for bacteria. Escaping from oxygen involves a clever response: whenever it finds a low oxygen environment it attaches to surfaces, associating into biofilms. Biofilms protect *S. epidermidis* against host cells. Understanding these responses is a must in order to develop treatments and prevent infection success.

## INTRODUCTION

In mammals, saprophytic microorganisms contribute to the control of pathogenic bacteria, digestion of nutrients and synthesis of diverse coenzymes, prosthetic groups and amino acids [1]. On the skin the microorganism population is estimated at 10^11^ to 10^12^ cells [2, 3]. In the dermic layers, the saprophytic bacterium *Staphylococcus epidermidis* inhibits colonization by *Staphylococcus aureus* or *Streptococcus pyrogenes* through secretion of proteases [4] and other antimicrobial compounds [5]. *S. epidermidis* is abundant in moist areas, occupying a micro-aerobic niche in dermis and epidermis and in the nearly anoxic sebaceous areas [6].

*S. epidermidis* may be accidentally internalized through wounds or punctures made after improper sterilizing procedures. Upon entry, this bacterium has to face high oxygen concentration [O_2_] and rejection from the immune system, a situation that most likely triggers a stress response [7]. As bacteria are carried through the organism, they may reach areas where low [O_2_], similar to that found in the skin layers that constitute its natural habitat, and it is likely that in an effort to remain in the hypoxic area, it adheres to epithelia or other surfaces and organizes into biofilms. This response protects *S. epidermidis* against both, phagocytosis and antibiotics [8]. In regard to low [O_2_] environments within the host, these are often found in the vicinity of artificial devices such as catheters or prosthetic valves, favoring bacterial biofilm formation and forcing implanted device removal [9, 10].

Understanding the *S. epidermidis* response to different [O_2_] would help optimize treatments [11]. Growing *S. epidermidis* at different [O_2_] leads to differential expression of redox enzymes in the respiratory chain and to different biofilm formation patterns [12]. At high [O_2_], high expression of cytochrome oxidases and NADH dehydrogenases is observed, while the tendency to form biofilms is minimal. In contrast, [O_2_] depletion increases nitrate reductase expression and biofilm generation [12].

The aerobic and anaerobic metabolism of *S. epidermidis* grown at different [O_2_] were evaluated. The rates of anaerobic fermentation and O_2_ consumption varied in opposite senses, i.e., increasing [O_2_] in the growth medium increased the rate of oxygen consumption and decreased the rate of fermentation and vice-versa: at low [O_2_], the rate of oxygen consumption decreased while fermentation increased. In addition, it was observed that at high [O_2_] the susceptibility of *S. epidermidis* to the toxic effects of hydrogen peroxide increased. Remarkably, in the micro- and anaerobic environments *S. epidermidis* growth was slower, while [ATP] was higher, probably indicating that cells were preparing to associate in biofilms [13]. When ATP synthesis was inhibited to different degrees by inhibitors of the respiration chain (cyanide) [12] or glycolysis (1, 4-bisphosphobutane) [14, 15], biofilm formation also decreased. For *S. epidermidis*, the advantage of anchoring itself in a biofilm would be to remain at its ideal [O_2_], while enhancing its resistance to antibodies or antibiotics [8]. To understand and prevent these processes, the physiology of *S. epidermidis* needs to be analyzed thoroughly.

## MATERIALS AND METHODS

### Bacterial strain and growth media

*S. epidermidis* strain ATCC 12228 was donated by Dr. Juan Carlos Cancino Díaz (Instituto Politécnico Nacional, México). A loophole from the bacterium was suspended in 5 mL of 3% tryptic soy broth (Fluka, Sigma) and incubated at 37ºC for 24 h. Subsequently, pre-cultures were added to 1 L LB medium (1% tryptone, 0.5% yeast extract, 1% NaCl) plus 2% glucose and incubated 24 hours at 30 ºC under aerobic (shaking 150 rpm), microaerobic (5% CO_2_, no agitation) or anaerobic (static in oxygen-depleted sealed acrylic chamber) conditions. Then the cells were washed three times at 5000 xg for 10 min with distilled water and resuspended in 10 mM HEPES pH 7.4

### Cytoplasmic extracts

All procedures were conducted at 4 °C. Cells (grown under aerobic, microaerobic or anaerobic conditions) were centrifuged at 5000 xg for 10 min, washed three times with distilled water and resuspended in 50 mL 10 mM HEPES, pH 7.4, supplemented with one tablet of protease-inhibitor cocktail (Complete) and 1 mM PMSF. Cells were disrupted by sonication using a Sonics VibraCell sonicator (Sonics & materials, Inc., Newtown, CT) 7 x 20 sec with 20 sec intervals. To remove unbroken cells the suspension was centrifuged at 10 000 xg for 10 min and the supernatant was recovered.

### Protein concentration

Protein concentrations from intact *S. epidermidis* cells were determined by the biuret method [16]. Absorbance at 540 nm was read in a Beckman-Coulter DU50 spectrophotometer. For cytoplasmic extracts, protein concentration was measured by Bradford at 595 nm, using 1 to 2 µL aliquots of the sample in a PolarStar Omega (BMG labtech, Ortenberg, Germany) [17].

### Rate of Oxygen Consumption

The rate of oxygen consumption was measured in a water-jacketed 1 mL chamber at 37 °C equipped with a Clark type electrode connected to a Strathkelvin model 782 oxymeter. Data were analyzed using the 782 Oxygen System Software (Warner/Strathkelvin Instruments) [12]. Reaction medium 10 mM HEPES pH 7.4 plus the indicated respiratory substrate: 33 mM ethanol, 10 mM lactate or 40 mM glucose. Bacteria, 0.5 mg prot mL^−1^.

### Ethanol production

Fermentation by cell cytoplasmic extracts (0.5 mg prot. mL^−1^) was measured in 0.1 M MES-TEA, pH 7.0, 1.8 mM NAD plus either 20 mM glucose or glycerol and incubated at 30 °C for 0, 2.5, 5 or 10 min. The reaction was stopped with 30% TCA, 0.1 mL and neutralized with NaOH. Ethanol was measured adding a 10 µL aliquot (0.005 mg) of the supernatant to 0.2 mL 114 mM K_2_HPO_4_, pH 7.6. After 1 min, 30 μg ADH mL^−1^ was added, the sample was incubated for 30 min and O.D. was determined at 340 nm in a POLARstar Omega. Ethanol is reported as μmol ethanol (mg prot)^−1^ [18].

### ATP concentration

The concentration of ATP was determined in cytoplasm extracts. Cytoplasmic extracts were resuspended to 0.025 mg protein in 0.15 mL reaction buffer (20 mM KH_2_PO_4_, 40 mM Na_2_HPO_4_, 80 mM NaCl plus 1 mM MgSO_4_). An ATP calibration curve was prepared freshly each day using lyophilized luciferase (Sigma-Aldrich). Luciferase was prepared following instructions by the provider and 0.02 mL was added to each sample in a 96-well microplate. Bioluminescence was detected in a POLARstar Omega luminometer (BGM LABTECH, Offenburg, Germany). [ATP] was reported as µmol (mg prot)^−1^ [19, 20].

### Susceptibility to Hydrogen peroxide-mediated damage

The effect of [H_2_O_2_] on the viability of *S. epidermidis* was determined as previously reported [21]. Briefly, cells grown under aerobiosis, microaerobiosis or anaerobiosis were adjusted to an O.D.= 0.1 (600 nm). Then H_2_O_2_ (0 to 25 mM as indicated) was added to the culture (final O.D.= 0.025). After 30 minutes, serial dilution of the cultures was performed in 0.9% NaCl. Afterwards 10 µL of the 1:1000 diluted sample was plated in LB, 2% glucose agar plates and incubated 24 h at 37 ºC. Colony forming units (CFU) were counted. CFU mL^−1^ before addition of H_2_O_2_ was assigned as 100%. The average of three experiments is shown with SD. ANOVA test and Tukey’s multiple comparison-test were used. Significance was \**P* < 0.0001.

### Biofilm formation and detection

The biofilm assay was performed using *S. epidermidis* cultures and sterile Costar 96-well polystyrene plates as previously reported with some modifications [12, 22]. Pre-cultures were grown overnight in trypticase soy broth (TSB) at 37°C. To evaluate the effect of different inhibitors, sodium cyanide (NaCN) 100 µM, butane-1,4-bisphosphate(B1,4BP) 1 mM, both inhibitors, carbonyl cyanide m-chlorophenyl hydrazone (CCCP) 6 µM and a control without treatment were added to the microplate at the beginning of the assay. Thereafter, the bacterial suspensions were diluted at D.O.= 0.02 and adjusted to 200 μL with fresh TSB. The plate was incubated 24 hours at 37 °C with 5% CO_2_. After incubation, the TSB medium was removed and the wells were washed twice with 200 µL of phosphate-buffered saline (PBS) to remove non-adherent bacteria. The plates were dried for 1 hour at 60 °C and stained with 0.4% crystal violet solution for 10 min. The plates were washed under running tap water to remove any excess stain. Biofilm formation was determined by the solubilization of the crystal violet stain in 200µL of 33% glacial acetic acid for 10 min, shaking and measuring the absorbance (492 nm) with a microplate reader (Polar Star Omega, BMG Labtech).. Each sample was tested in three independent experiments in triplicate and compared with the control (without treatment) using one-way analysis of variance (ANOVA) followed by Dunnett’s post hoc test.

### Genome analysis and Protein Modeling

The genomic sequence of *S. epidermidis* ATCC 12228 was obtained from the NCBI database (GenBank: CP022247.1) [23]. NADH DH structures (NCBI protein id: ASJ93946.1, ASJ94976.1, ASJ93963.1) were modeled using 3D homology models from the Swiss Model- Expasy, Swiss Inst. of Bioinformatics Biozentrum, Univ. of Basel https://swissmodel.expasy.org

## RESULTS

Oxygen is among the most important factors driving evolution. Its partial reduction products, the reactive oxygen species (ROS) destroy nucleic acids, proteins and membranes [24]. Thus, in spite of its remarkable electron acceptor properties, the dangerous oxygen molecule has to be dealt with carefully [25, 26]. *S. epidermidis* lives in hypoxic/anoxic environments, although it can adapt to aerobic environments. In order to follow the adaptation of *S. epidermidis*, it was cultivated at different [O_2_]. Cells grew proportionally to [O_2_]. That is, under aerobic conditions biomass yield was 8.58 g(ww)/L, three times higher than under microaerobiosis where biomass was 2.11 g(ww)/L or under anaerobiosis at 1.75 g(ww)/L.

Biomass data evidenced that at high [O_2_] cells grew better. In order to further explore the basis for the large increase in biomass under aerobic conditions, it was decided to test the activity of the respiratory chain from *S. epidermidis* grown at different [O_2_] (Figure 1). As expected, the ability of cells to consume oxygen was proportional to [O_2_] in the growing environment, demonstrating the adaptability of *S. epidermidis* to varying conditions. In aerobic conditions, the rate of oxygen consumption in the presence of lactate was 70 natgO (mg prot.min)^−1^, at least five times higher than in microaerobic media, where the rate was 5 natgO (mg prot.min)^−1^ or in those grown under anaerobic conditions, where it became negligible (Figure 1). In addition, it was observed that under aerobic conditions the best respiratory fuel was lactate, which was consumed around three times as fast as glucose or ethanol (Figure 1).

**Figure. 1.**
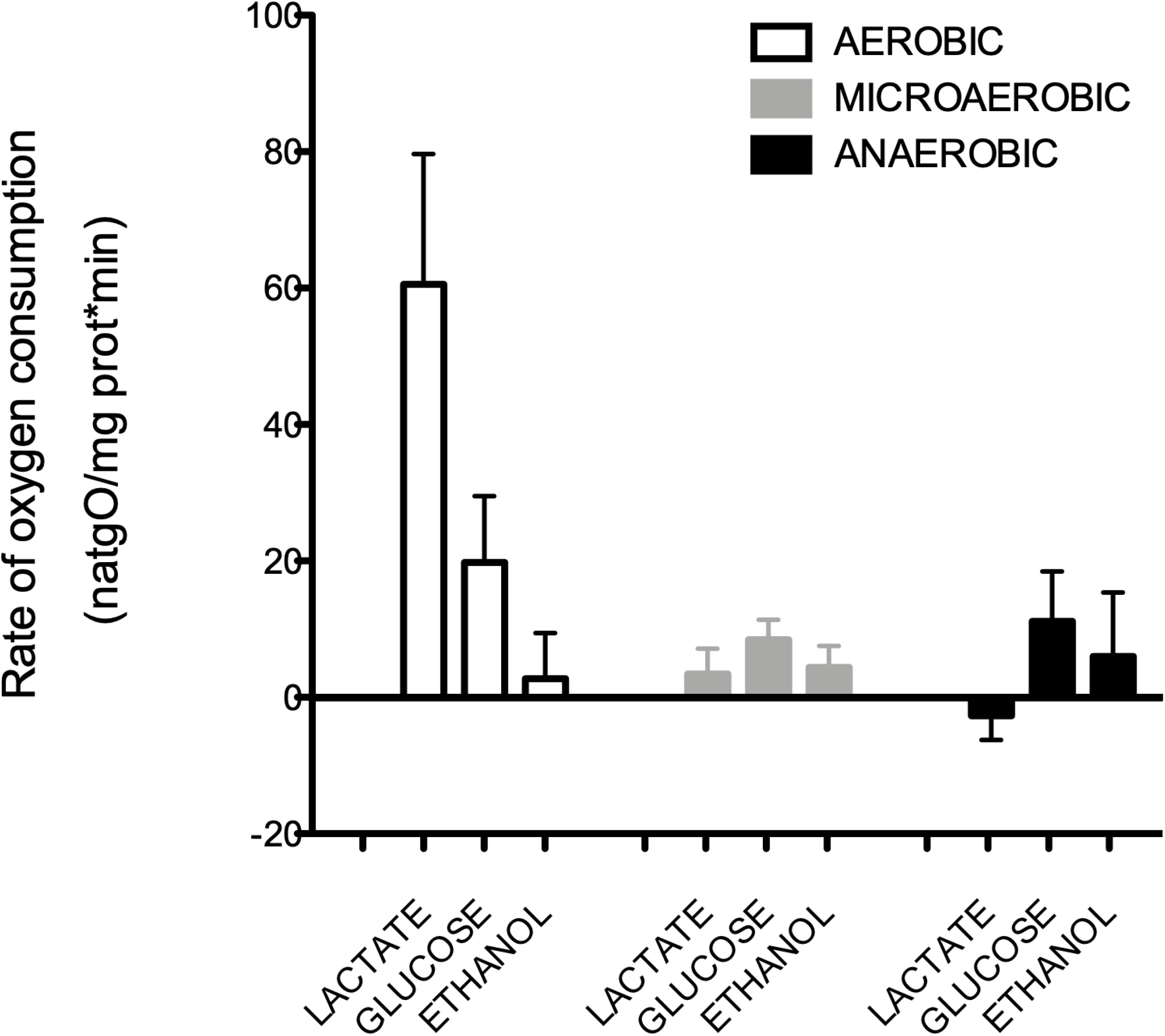
Rate of oxygen consumption by *S. epidermidis* in the presence of different respiratory substrates. Experimental conditions: 10 mM HEPES (pH 7.4), 10 mM lactate, 40 mM glucose or 33 mM ethanol. Cells were grown at different [O_2_] as follows: aerobic (empty bars), microaerobic (gray bars) and anaerobic (black bars).

The activity of the respiratory chain correlated with the different growth rates of *S. epidermidis*. However, glycolysis can also constitute an important source of energy [27]. In fact, as low oxygen environments seem to be preferred by *S. epidermidis*, anaerobic glycolysis should be the preferred energy-yielding pathway in this bacterium. To test this, *S. epidermidis* was grown in the presence of different [O_2_] and its ability to perform anaerobic glycolysis was measured. Substrates were glucose (Figure 2-A) or glycerol (Figure 2-B). The efficiency of *S. epidermidis* to ferment these two substrates was roughly equivalent. Also, measurements made at 2.5, 5 and 10 min of incubation did not result in modifications in the concentration of ethanol. When grown at different [O_2_], bacteria from anaerobic media were the most active, fermenting both glucose and glycerol (Figure 2), suggesting that glycolytic activity changes in the opposite sense as [O_2_].

**Figure. 2.**
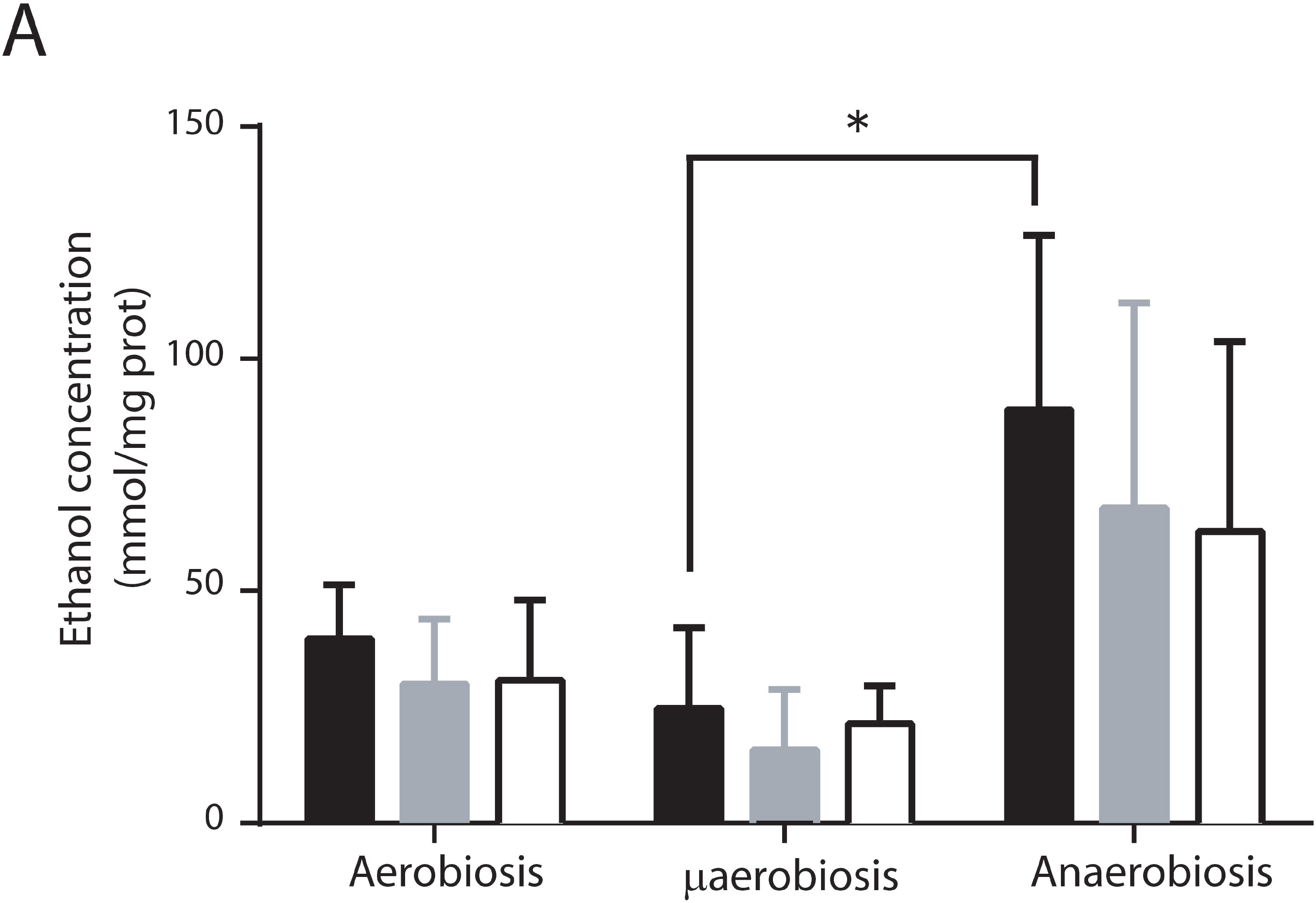
Fermentation by *S. epidermidis* grown at different [O_2_]. Cytoplasmic extracts were obtained from *S. epidermidis* grown under aerobic, microaerobic or anaerobic conditions. Fermentation by cell cytoplasmic extracts (0.5 mg prot. mL^−1^) was measured using A) 20 mM glucose or B) 20 mM glycerol. Samples were incubated at 30 °C for: 2.5 min (black columns), 5 min (gray columns) or 10 min (white columns). Results are reported as μmol ethanol per mg protein. Tukey’s comparison test was used to determine significant differences (\**P* < 0.05).

Activation of glycolysis under anaerobiosis evidenced that not only the aerobic, but also the anaerobic metabolism of the cell adapts to [O_2_] in the culture medium. Thus, we decided to measure the effect that growing the cells at different [O_2_] would have on the concentration of ATP ([ATP]) (Figure 3). Contrary to what we expected from the low respiratory activity plus the small increase in fermentation activity observed under micro- and anaerobiosis, [ATP] increased at low [O_2_]. The increase in [ATP] was roughly five times in microaerobiosis and three times in anaerobiosis (Figure 3). Aiming to understand such rise in ATP, we found that other bacteria, e.g. *Bacillus brevis* and *Escherichia coli*, react to substrate depletion by adhering to glass surfaces and at the same time increase [ATP] two-to fivefold in comparison to planktonic cells [28]. In this regard, it has been reported that hypoxic stimuli induce biofilm formation in *S. epidermidis* [12].

**Figure 3.**
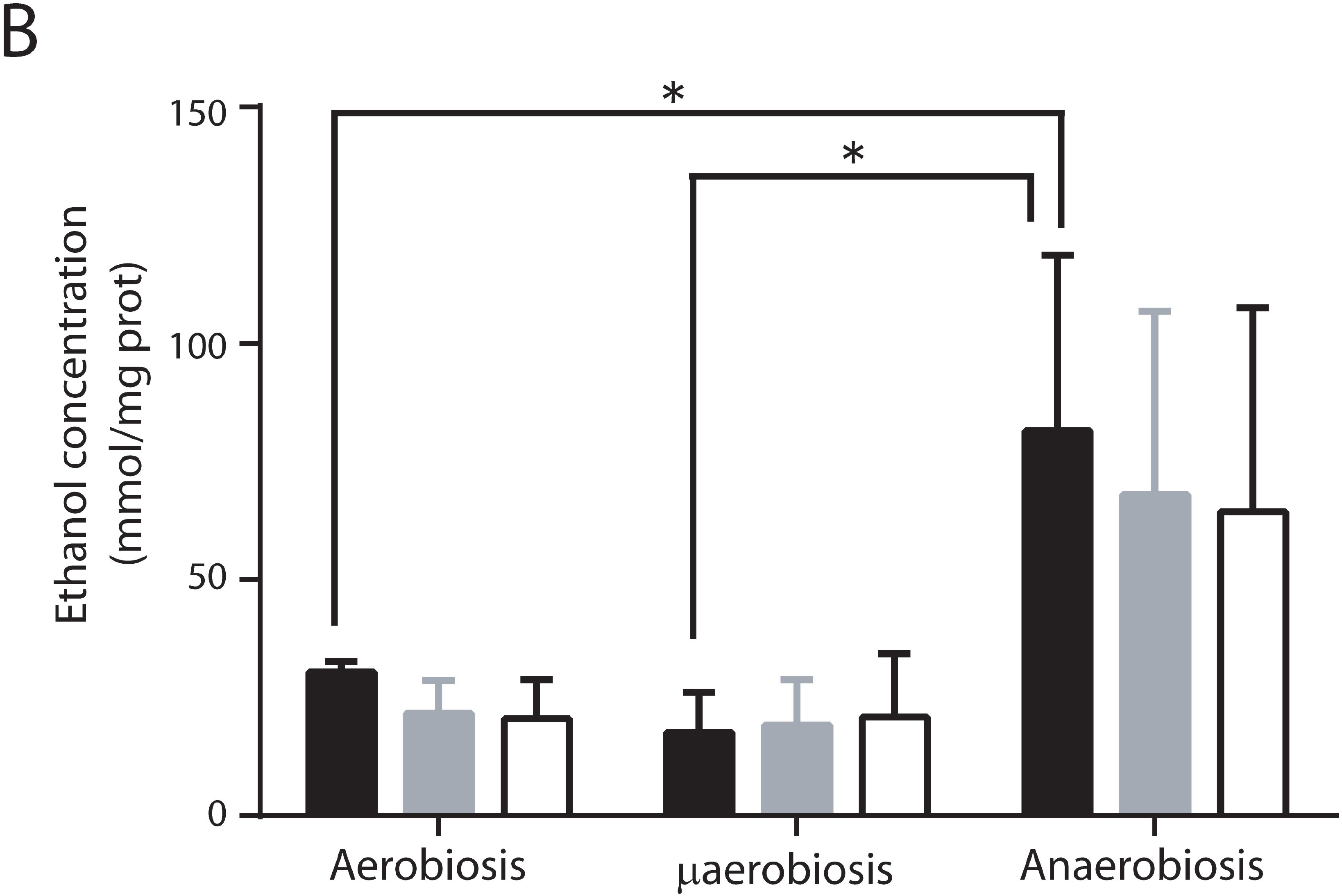
Intracellular ATP concentrations in *S. epidermidis* grown at different [O_2_]. Cells were grown at different [O_2_] in LB plus glucose. Cytoplasmic extracts were obtained from each of these cultures and used to measure intracellular ATP. ATP concentration was estimated using luciferase and interpolating into a standard curve (See methods). The average of three experiments is shown with SD. * indicates significant difference *P* < 0.05.

In *Staphylococcus aureus*, expressing a deficient respiratory chain decreases sensitivity to H_2_O_2_ [29], suggesting that anaerobiosis-adapted cells resist oxidative stress better. Here, as oxygen decreased in the growth medium, *S. epidermidis* switched its metabolic mode from aerobic to fermentative. Thus, we decided to test the sensitivity to ROS of *S. epidermidis* grown at different [O_2_] to the presence of hydrogen peroxide (Figure 4) [30]. Even at the lowest concentrations of H_2_O_2_ we used (0.5 mM), viability decreased in all cells. However, in all cases the aerobic-grown cells exhibited the poorest survival rates, while cells grown under anaerobiosis survived H_2_O_2_ best, such that even at the highest H_2_O_2_ concentration tested (25 mM H_2_O_2_) a small amount of viable cells was detected (Figure 4). The increase in sensitivity to ROS observed in aerobically grown *S. epidermidis* was probably due to increased expression of the redox enzymes in the respiratory chain [12]. These redox enzymes contain different coenzymes and prosthetic groups, which normally become free radicals during their catalytic cycle that are perhaps activated by H_2_O_2_ [26, 31].

**Figure 4.**
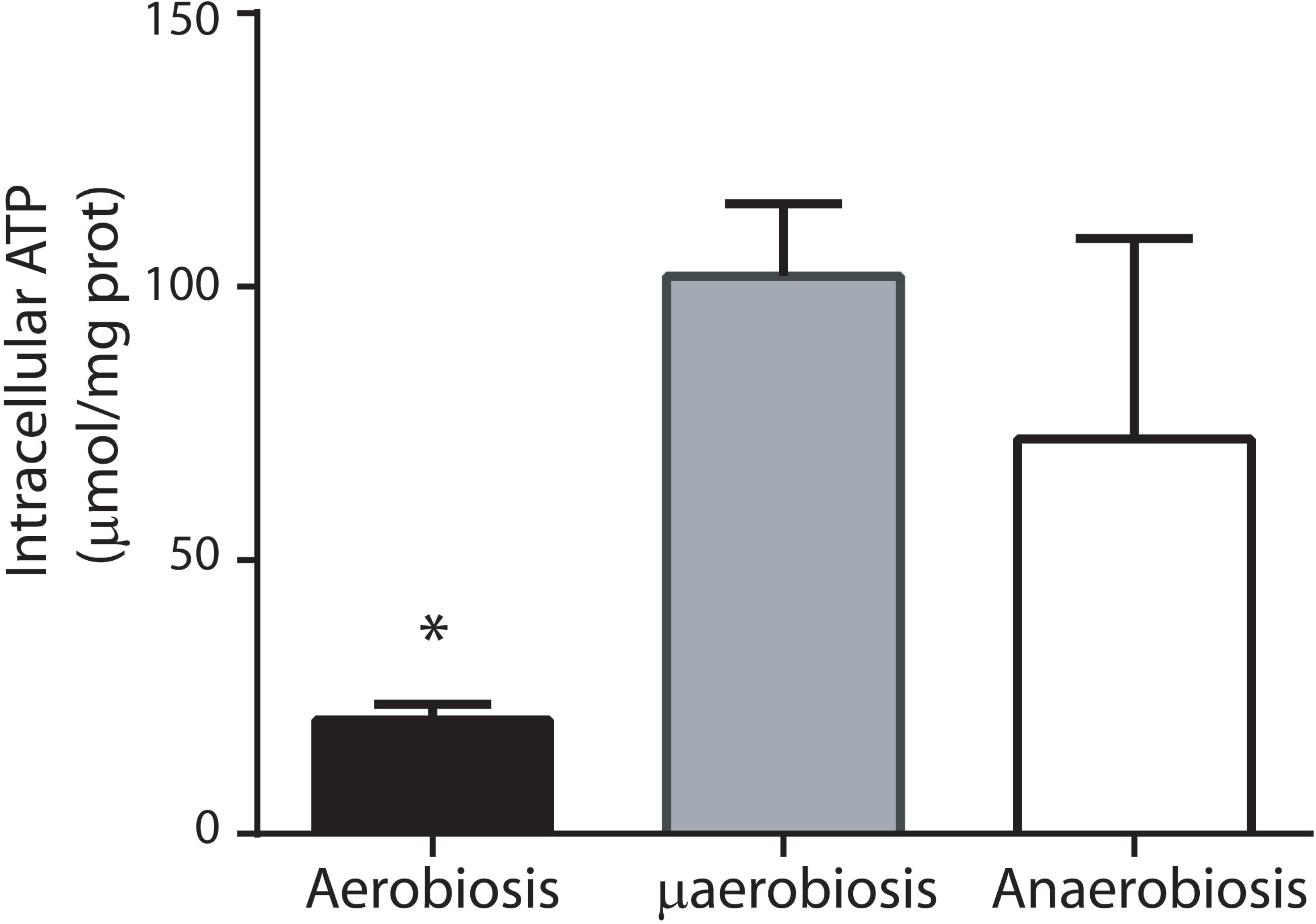
H_2_O_2_ effect on cellular viability. *S. epidermidis* susceptibility to hydrogen peroxide was determined using 0, 0.5, 1, 5, 10 or 25 mM of H_2_O_2_ in each group: aerobiosis (black bar), microaerobiosis (gray bar) or anaerobiosis (white bar). After 30 min of incubation with H_2_O_2,_ the samples were diluted 1:1000, 10 µl were taken and plated in LB plus 2% glucose-agar. CFU/mL were counted taking the samples without treatment as 100% viable cells. The average of three experiments is shown with SD. Significance \**P* < 0.0001.

When grown at high [O_2_] *S. epidermidis* expressed an active respiratory chain. However, adaptation to high [O_2_] also resulted in increased sensitivity to hydrogen peroxide. This probably indicates that there is a trade-off between ROS protection and higher growth in *S. epidermidis* according to the oxygen concentration. Which is why even when [ATP] was higher in cells grown at low [O_2_], their rate of growth decreased. This seemingly contradictory situation may be explained by proposing that when *S. epidermidis* finds a low [O_2_], which resembles that found in its normal niche, it redirects its ATP from growth to production of polysaccharides and proteins involved in biofilm generation [13, 32]. To analyze the ATP-dependence of biofilm formation, the ability of cells grown in hypoxic conditions to form biofilms was analyzed. It was observed that cells incubated in the presence of the respiratory chain inhibitor cyanide or the glycolytic inhibitor 1,4-bisphosphobutane, formed smaller biofilms than the control and that addition of both inhibitors led to even smaller biofilms (Figure 5). This would suggest that biofilm formation activity is proportional to [ATP]. Complete ATP depletion by the uncoupler (CCCP) could not be evaluated due to its toxicity, which kills cells at 6 µM (Result not Shown).

**Figure 5.**
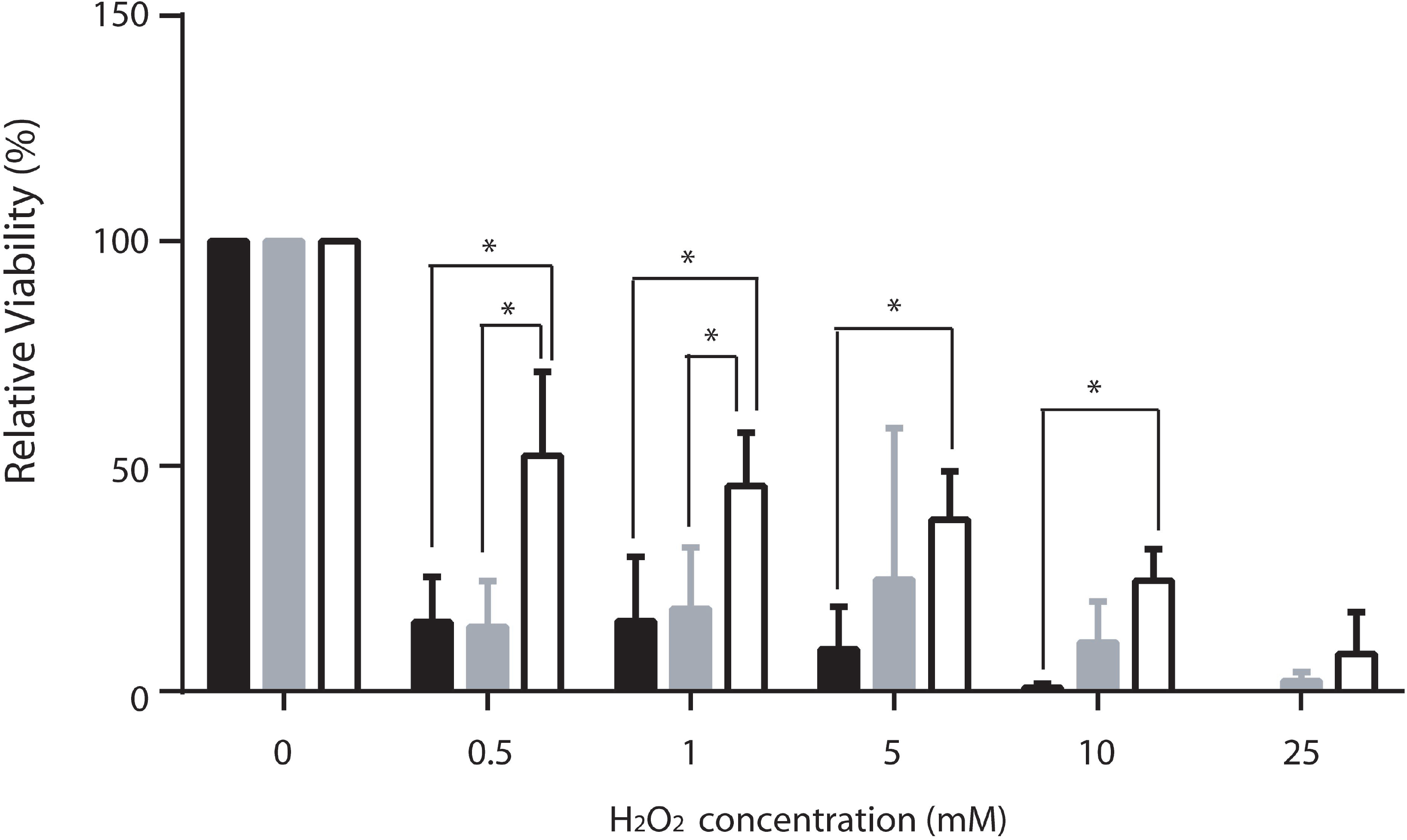
*In vitro* biofilm inhibition assay. *S. epidermidis* in microaerobic condition biofilm formation was determined using 100 µM NaCN, 1 mM B1,4BP or both inhibitors and compared to a control without inhibitors. After 24 hours of incubation, biofilm generation was evaluated by measuring the absorbance to 492 nm with a microplate reader. Each sample was compared with the control (without treatment) using ANOVA and Dunnett’s post hoc test. Significance \**P* < 0.0001.

## DISCUSSION

Diverse facultative bacteria, adapt to the [O_2_] in the medium, differentially expressing redox enzymes in its respiratory chain. *S. epidermidis* does express different enzymes at varying [O_2_][12]. This traduced in different rates of oxygen consumption and growth. Aerobic metabolism enabled cells to grow more [33]. Still, enhanced growth resulted in higher sensitivity to H_2_O_2_, suggesting that high contents of redox enzymes make cells vulnerable to ROS. Indeed, when grown at high [O_2_], ROS sensitivity of *S. aureus* and *Enterococcus faecalis* increases, while their mutant counterparts, lacking an efficient respiratory chain resist ROS better [29].

When exposing *S. epidermidis* grown in different [O_2_] to oxygen peroxide, we observed a similar phenomenon: cells grown in hypoxic or anoxic environments, which exhibited low respiratory rates were more resistant to oxygen peroxide (Figure 4). Thus, as in *S. aureus*, the lack of an efficient respiratory chain in *S. epidermidis* enabled cells to survive ROS. This is probably useful when bacteria are confronted with the oxidative burst generated by the immune system.

The rate of oxygen consumption in aerobic grown cells was highest when lactate was the substrate. This is probably due to the direct donation of electrons to the menaquinone pool by lactate dehydrogenase [34, 35]. The slower rates observed for alcohol, may be due to an additional step as alcohol dehydrogenase electrons are first donated to Ndi2 [36]. The rate of respiration was also slow for glucose, probably for the same reason, that its intermediaries have to undergo many reactions before releasing electrons to the respiratory chain [37]. In contrast, under anaerobiosis, lactate-dependent oxygen consumption disappeared completely while a small rate of glucose-dependent oxygen consumption was still present. In contrast, in *S. aureus* increased lactate dehydrogenase expression anaerobiosis has been reported [38]

In addition to lactate dehydrogenase, among the differentially expressed redox enzymes in both *S. epidermidis* [23] and *S. aureus* are Ndi2s, substituting complex I as in common in facultative bacteria. Structural modeling isoform 1 (Suppl Fig. 1A) and isoform 2 (Suppl. Fig. 1B) illustrated the similarities of these enzymes to their counterparts in *S. aureus* [39]. In addition, *S. epidermidis* isoform three (Suppl. Fig 1C) was modeled successfully using the internal NADH dehydrogenase Ndi2 from the facultative yeast *S. cerevisiae* [40]. Thus, it is likely that Ndi2s may be used to pred(ict the facultative nature of a species.

The normal habitat for *S. epidermidis* is the microaerobic environment found in different epidermic and dermic layers [41]. One strategy *S. epidermidis* uses when confronted with high [O_2_] is the differential expression of a diverse number of redox enzymes in the respiratory chain. Reports indicate that when microaerophilic or anaerophilic bacteria find a suitable environment, they react manufacturing proteins and polysaccharides that enable them to form biofilms and attach to surfaces at low [O_2_]. Avoiding high [O_2_] involves both, anchoring in low oxygen environments and building biofilms as barriers against penetration of ROS or toxic substances [20]. Metabolic adaptation has also been reported for *Neisseria gonorrhoeae*, when it is stimulated to form biofilms. A proteomic analysis of *N. gonorrhoeae* biofilms evidenced up-regulation of proteins involved in anaerobic metabolism such as glycolysis and TCA cycle plus increased expression of those proteins involved in biofilm generation like pilus-associated proteins [42]. In addition, some oxidative stress genes are required for normal biofilm formation in *N. gonorrhoeae* [43].

The increase in ATP prior to biofilm formation has been reported in other bacteria. *Bacillus brevis and Escherichia coli* react to substrate depletion by adhering to glass surfaces and at the same time increase [ATP] two-to fivefold as compared to planktonic cells [28]. So the conditions where bacteria need to make biofilms promote saving ATP even at the expense of the growth rate. ATP is most likely needed to synthesize the extracellular proteins and the polysaccharide fibers that anchor cells to surfaces and to each other. Inhibiting ATP production in micro- or anaerobic conditions by adding cyanide or 1,4-bisphosphobutane resulted in a reduced biofilm formation (Figure 5). This phenomenon is also observed when treating *S. epidermidis* with the nitrate reductase inhibitor methylamine in anaerobic conditions [12]. In contrast, in aerobiosis cyanide promotes biolfilm formation [12].

Even when facultative bacteria such as *S. epidermidis* survive at high [O_2_], their habitat in the skin is hypoxic to anoxic. While they survive in aerobic environments their susceptibility to ROS-mediated damage and possibly to attack by macrophages increases. They thus present an oxygen avoidance behavior, anchoring and associating in hypoxic environments (Figure 6). Learning how avoidance works in *S. epidermidis* and other bacteria would impact both the physiologic and therapeutic field.

**Figure 6.**
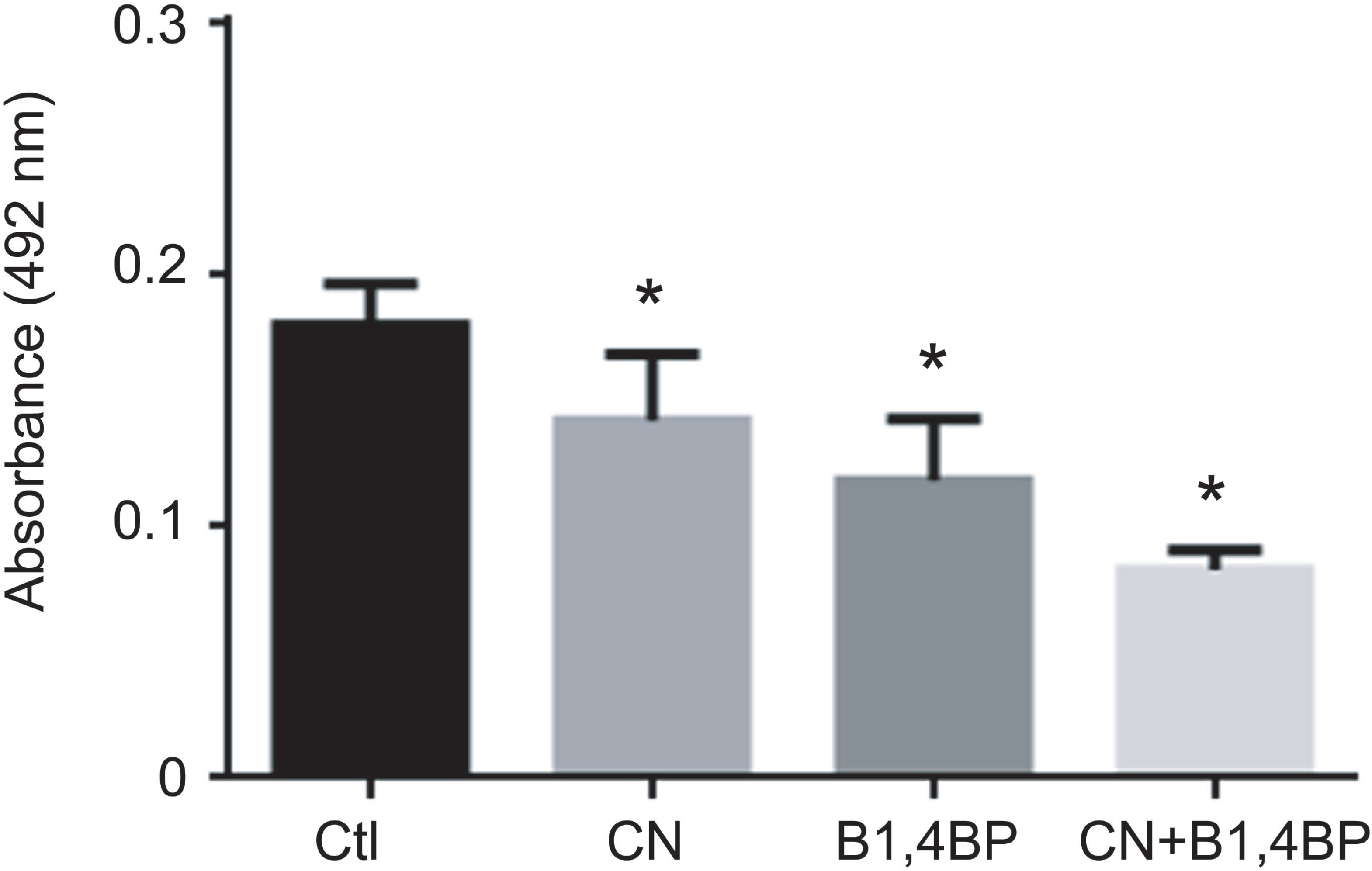
Cartoon depicting the shift that *Staphylococcus epidermidis* makes when [O_2_] decreases in the growth medium. (A). When high oxygen concentrations are found in the medium, *S. epidermidis* cells are soluble, as planktonic cells and may flow with the blood in vessels. (B) In contrast, under micro- or anaerobic conditions the cells shift to a fermentative metabolism and ATP accumulates in preparation for adhesion to a suitable surface (i.e. epithelia, catheters, valves) and the eventual formation of a biofilm. In this state the cells exhibit more resistance to H_2_O_2_ mediated damage. Excess ATP is probably used to produce adhesion proteins and poly-N-acetylglucosamine (gray fibers in the illustration)

## Acknowledgements

Partially funded by UNAM/DGAPA/PAPIIT IN203018. UHPD (MsSc) and EES (PhD) are graduate students in the Biochemistry Program at UNAM. LMG is in the Biomedical PhD program at UNAM. UHPD and LMG are CONACYT fellows. CUA present address: Fox-Chase Cancer Center, Philadelphia, PA. Technical help from Ramón Méndez-Franco is acknowledged. Authors do not have any conflicts of interests.

---

**Figure S1.**
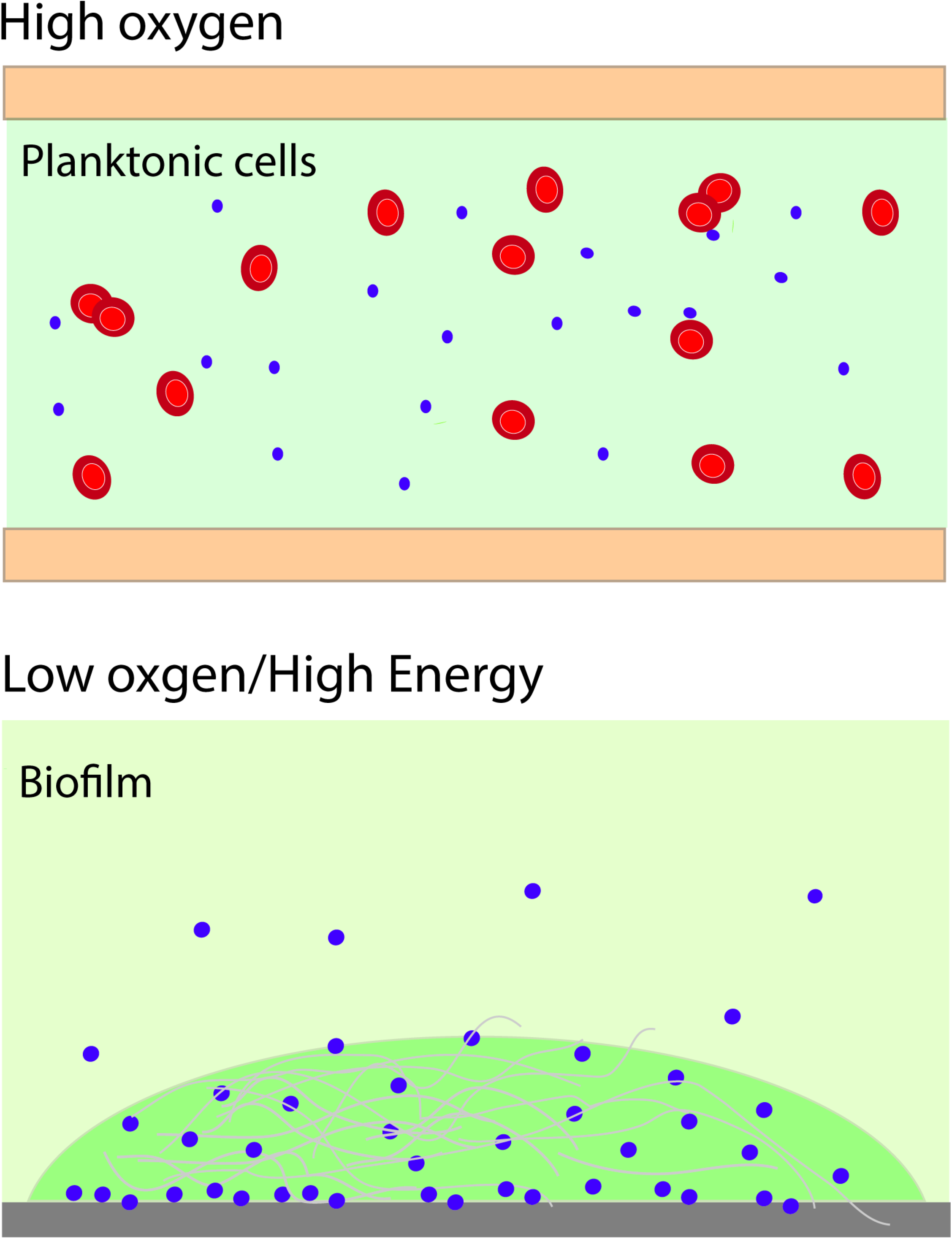
Modeling of NADH type II dehydrogenases from *S. epidermidis*. Images on the left are models made after the templates shown on the right. (a) *S. epidermidis* NCBI protein id: ASJ93946.1 modeled after the crystal structure of a putative NADH-dependent flavin oxidoreductase from *S. aureus* SMTL ID: 3I5a.1 (Lam, R. et al. unpublished). (b) *S. epidermidis* NCBI protein id: ASJ94976.1 modeled using the crystal structure of a NADH-quinone oxidoreductase (NDH-II) from *S. aureus* E172S mutant SMTL ID: 5na4.1 [34]. (c) S. *epidermidis* NCBI protein id: ASJ93963.1 modeled using as a template the crystal structure of the Ndi1 protein from *Saccharomyces cerevisiae* in complex with the competitive inhibitor stigmatellin SMTL ID: 5yjw.1 [40].

